## Supplementary Data and Figures for "The Helix-Loop-Helix motif of human EIF3A regulates translation of proliferative cellular mRNAs"

#### Supplementary Tables

Included in a single Excel .xlsx file:
Table S1. RNAseq data (Supplementary Tables, tab 1, “RNAseq nonzero”)
Table S2. Ribosome profiling data (Supplementary Tables, tab 2, “RP (RSnonzero match)”)
Table S3. Babel statistical analysis of RNAseq and ribosome profiling data (Supplementary Tables, tab 3, “Babel_TE_FC”)
Table S4. Statistically significant transcripts from Babel analysis (Supplementary Tables, tab 4, “Babel_p<0.01”)
Table S5. Functional classification of translationally downregulated hits with PMID (Supplementary Tables, tab 4, “S5 Babel_p<0.01 FC0.4”)
Table S6. Functional classification of translationally upregulated hits with PMID (Supplementary Tables, tab 4, “S5 Babel_p<0.01 FC2.5”)

#### Cloning

EIF3A was PCR-amplified from HEK293T cDNA and cloned into vector nLv-103 (hygromycin) using PCR-based restriction free cloning. HLH mutations and shRNA target site mutations were introduced by PCR-based site-directed mutagenesis. Custom EIF3A shRNA lentiviral vector was obtained from Sigma-Aldrich MISSION TRC shRNA pLKO.1-puro bacterial stock (Sigma SHCLNG). pMD2.G (Addgene #12259) envelope protein vector and pCMV-dR8.91 (Addgene # 2221) packaging vector were used to make viral particles for transduction. The same vectors were later used to package CRISPRi lentiviral plasmids. HCV IRES, *MYC* 5’-UTR, and *ATF4* 5’-UTR elements were cloned into pcDNA4-Rluc using HinDIII and EcoRI restriction sites; the *HBB* 5’-UTR was cloned into pcDNA4-Fluc, where the *Renilla* luciferase ORF sequence was replaced by the *Firefly* luciferase ORF. *ATF4* 5’UTR uORF1 and uORF2 mutations were introduced by PCR-based site-directed mutagenesis. pCMV-SPORT6 containing MGC Human *MYC* cDNA (CloneId:2985844) glycerol stock was obtained from Dharmacon (MHS6278). Full-length *MYC* and *GAPDH* were amplified from cDNA and cloned into pCMV-SPORT6 using SalI and NotI restriction sites. pHR-EF1a-dCas9-HA-BFP-KRAB-NLS, a gift from the Jacob Corn Lab (Addgene plasmid #102244), was used to introduce catalytically dead Cas9 into HEK293T cell lines. EIF3A sgRNA sequences were obtained from the human CRISPRi library v2[20] and cloned into in pSLQ1371_BLP1_Ef1A_puro_GFP, a gift from the Jonathan Weissman Lab, using BstX1 and Blp1 restriction sites. The EIF3A sgRNA top and bottom oligo were ordered from IDT (for final selected sgRNA: (top) 5’- TTGGCAGCCGGCCAGAGACGGAAGTTTAAGAGC -3’; (bottom) 5’- TTAGCTCTTAAACTTCCGTCTCTGGCCGGCTGCCAACAAG -3’) and annealed by incubating at 95 ºC for 5 min in annealing buffer (100 mM Potassium acetate, 30 mM HEPES-KOH pH 7.4, 2 mM Mg acetate) and slowly cooling to room temperature. 5 nM annealed oligo were ligated to 10 ng digested vector backbone using T4 ligase (Thermo A13726) according to the manufacturer’s protocol.

*EIF3A* mRNA (site of HLH mutations indicated in red)
ATGCCGGCCTATTTTCAGAGGCCGGAAAATGCCCTCAAACGCGCCAACGAATTTCTTGAGAA
CATAATTGGTTCGTGTATCTTTTGCCATGTTCTTTATGATGTTATGAAAAGTAAAAAACATAG
AACATGGCAAAAGATACACGAACCAATTATGTTGAAATACTTGGAACTTTGCGTGGATCTTC
GCAAGAGCCACTTGGCAAAGGAGGGGTTATACCAGTATAAGAACATTTGTCAACAGGTGAA
CATAAAATCTCTGGAGGATGTTGTTAGGGCATATTTGAAAATGGCAGAGGAAAAAACTGAA
GCTGCTAAAGAAGAATCTCAGCAGATGGTCTTAGATATAGAGGATCTAGATAATATTCAAAC
TCCTGAGAGTGTTCTCCTAAGTGCTGTAAGTGGTGAAGACACTCAGGATCGTACTGACAGAT
TACTTTTAACTCCATGGGTTAAATTCCTGTGGGAGTCTTACAGGCAGTGTTTGGACCTTCTTA
GAAACAATTCTAGAGTAGAGCGCCTGTACCATGATATTGCCCAGCAAGCTTTCAAATTCTGC

CTCCAATACACGCGTAAGGCTGAATTCCGTAAACTGTGTGACAATTTGAGAATGCACTTATC
GCAGATTCAGCGCCACCATAACCAAAGTACGGCAATCAATCTTAATAATCCAGAGAGCCAG
TCCATGCATTTGGAAACCAGACTTGTTCAGCTGGACAGTGCTATCAGCATGGAATTGTGGCA
GGAAGCATTCAAAGCTGTGGAAGATATTCACGGGCTATTCTCCTTGTCTAAAAAACCACCTA
AACCTCAGTTGATGGCAAATTACTATAACAAAGTCTCAACTGTGTTTTGGAAATCTGGAAAT
GCTCTTTTTCATGCATCTACACTCCATCGTCTTTACCATCTCTCTAGAGAAATGAGAAAGAAT
CTCACACAAGATGAGATGCAAAGAATGTCTACTAGAGTCCTTTTAGCCACTCTTTCCATCCCT
ATTACTCCTGAGCGTACGGATATTGCTCGACTTCTGGATATGGATGGCATTATAGTTGAAAA
ACAGCGTCGCCTTGCAACACTACTAGGTCTTCAAGCCCCACCGACACGAATTGGCCTTATTA
ATGATATGGTCAGATTTAATGTACTACAATATGTTGTCCCAGAAGTGAAAGACCTTTACAAT
TGGCTTGAAGTAGAATTTAACCCATTAAAACTCTGTGAGCGAGTCACAAAGGTTCTAAATTG
GGTTAGGGAACAACCTGAAAAGGAACCGGAATTGCAGCAGTATGTGCCACAACTGCAAAAC
AACACCATCCTCCGCCTTCTGCAGCAGGTGTCACAGATTTATCAGAGCATTGAGTTTTCTCGT
TTGACTTCTTTGGTTCCTTTTGTTGATGCTTTCCAACTGGAACGGGCCATAGTAGATGCAGCC
AGGCATTGCGACTTGCAGGTTCGTATTGATCACACTTCTCGGACCCTGAGTTTTGGATCTGAT
TTGAATTATGCTACTCGAGAAGATGCTCCGATTGGTCCTCATTTGCAAAGCATGCCTTCAGA
GCAGATAAGAAACCAGCTGACAGCCATGTCCTCAGTACTTGCAAAAGCACTTGAAGTCATTA
AACCAGCTCATATACTGCAAGAGAAAGAAGAACAGCATCAGTTGGCTGTCACTGCATACCTT
AAAAATTCACGAAAAGAGCACCAGCGGATCCTGGCTCGCCGCCAGACAATTGAGGAGAGAA
AAGAGCGCCTTGAGAGTCTGAATATTCAGCGTGAGAAAGAAGAATTGGAACAGAGGGAAGC
TGAACTCCAGAAAGTGCGGAAGGCTGAGGAAGAGAGGCTGCGCCAGGAAGCAAAGGAGAG
AGAGAAGGAGCGTATCTTACAGGAACATGAACAAATCAAAAAGAAAACTGTCCGAGAGCGT
TTGGAGCAGATCAAGAAAACAGAACTGGGTGCCAAAGCATTCAAAGATATTGATATTGAAG
ACCTTGAGGAATTGGATCCAGATTTTATCATGGCTAAACAGGTTGAACAACTGGAGAAAGA
AAAGAAAGAACTTCAAGAACGCCTAAAGAATCAAGAAAAGAAGATTGACTATTTTGAAAGA
GCCAAACGTTTGGAAGAAATTCCTTTGATAAAGAGCGCTTACGAGGAACAGAGAATTAAAG
ACATGGATCTGTGGGAGCAACAAGAGGAAGAAAGAATTACTACAATGCAGCTAGAACGTGA
AAAGGCTCTTGAACATAAGAATCGAATGTCACGAATGCTTGAAGACAGAGATTTATTCGTAA
TGCGACTCAAAGCTGCACGGCAGTCTGTTTATGAGGAAAAACTTAAACAGTTTGAAGAGCG
ATTAGCAGAAGAAAGGCATAATCGATTGGAAGAACGGAAAAGGCAGCGTAAAGAAGAACG
CAGGATAACATACTATAGAGAAAAAGAAGAGGAGGAGCAGAGAAGGGCAGAAGAACAAAT
GCTAAAAGAGCGGGAAGAGAGAGAGCGCGCCGAACGAGCAAAACGCGAGGAAGAGCTACG
AGAGTATCAGGAGCGGGTGAAGAAATTAGAAGAAGTGGAAAGGAAAAAACGCCAAAGGGA
GTTGGAAATTGAAGAACGAGAACGGCGTAGAGAGGAAGAGAGAAGACTTGGCGATAGTTC
CCTTTCTAGAAAGGACTCTCGTTGGGGAGATAGAGATTCAGAAGGCACCTGGAGAAAAGGA
CCTGAAGCAGATTCTGAGTGGAGAAGAGGCCCGCCAGAGAAGGAGTGGAGACGTGGAGAA
GGGCGAGATGAGGACAGGTCTCATAGAAGAGATGAAGAGCGGCCCCGGCGTCTGGGGGAT
GATGAAGATAGAGAGCCCTCTCTTAGACCAGACGATGATCGGGTTCCCCGGCGTGGCATGG
ATGATGACAGAGGCCCTAGACGTGGTCCTGAGGAAGATAGGTTCTCTCGTCGTGGGGCAGA
CGATGACCGGCCTTCCTGGCGTAACACAGATGATGACAGGCCTCCCAGACGAATTGCCGATG
AAGACAGGGGAAACTGGCGTCATGCGGATGATGACAGACCACCTAGACGAGGACTGGATGA
GGACAGAGGAAGCTGGCGAACAGCTGATGAGGACAGAGGACCAAGACGTGGGATGGATGA
TGACCGGGGGCCGAGGCGAGGAGGCGCTGATGATGAGCGATCATCCTGGCGTAATGCTGAT
GATGACCGGGGTCCCAGGCGAGGGTTGGATGATGATCGGGGTCCCAGGCGAGGCATGGATG
ATGACCGGGGTCCCAGGCGAGGCATGGATGATGACCGGGGTCCCAGGCGAGGCATGGATGA
TGACCGGGGTCCCAGGCGAGGGTTGGATGATGATCGAGGACCTTGGAGGAACGCCGATGAT
GACAGAATTCCCAGGCGTGGTGCAGAGGATGACAGGGGCCCTTGGAGAAACATGGATGATG
ATCGCCTTTCAAGACGTGCTGATGATGATCGGTTTCCCAGACGGGGTGATGACTCAAGACCT
GGTCCTTGGAGACCATTAGTCAAGCCAGGTGGATGGAGAGAGAAAGAAAAAGCCAGAGAG
GAGAGCTGGGGTCCACCTCGAGAATCAAGGCCATCAGAAGAACGTGAATGGGACAGAGAA
AAAGAAAGGGACAGAGATAATCAAGATCGGGAGGAGAATGACAAGGACCCTGAGAGAGAA

AGGGACAGAGAGAGAGATGTGGATCGAGAGGATCGCTTCAGAAGACCTAGGGATGAAGGT
GGCTGGAGAAGAGGACCAGCTGAGGAATCTTCAAGCTGGAGAGACTCAAGTCGCCGGGACG
ATAGGGATAGGGATGACCGTCGCCGTGAGAGGGATGACCGGCGTGATCTAAGAGAAAGACG
AGATCTAAGAGACGACAGGGACCGAAGAGGACCTCCACTCAGATCAGAACGTGAAGAAGT
AAGTTCTTGGAGACGTGCTGATGACAGGAAAGATGACCGGGTGGAAGAGCGGGACCCTCCT
CGTCGAGTTCCTCCCCCAGCTCTTTCAAGAGACCGAGAAAGAGACCGAGACCGAGAAAGAG
AAGGTGAAAAAGAGAAGGCCTCATGGAGAGCTGAGAAAGATAGGGAATCTCTCCGTCGTAC
TAAAAATGAGACTGATGAAGATGGATGGACCACAGTACGACGTTAG

*EIF3A* HLH* - AAAAGTAAAAA mutated to AACAGTGAAGA
*EIF3A* shRNA target – GCGCCTTGAGAGTCTGAATAT
*EIF3A* shRNA target (mutated) – GCGACTAGAAAGCCTAAACAT
*EIF3A* HLH* fasta file sequence (for building Bowtie2 indices) –
ATGTTCTTTATGATGTTATGAACAGTGAAGACATAGAACATGGCAAAAG

HCV IRES (start codon indicated in red)
CTCCCCTGTGAGGAACTACTGTCTTCACGCAGAAAGCGTCTAGCCATGGCGTTAGTATGAGT
GTCGTGCAGCCTCCAGGACCCCCCCTCCCGGGAGAGCCATAGTGGTCTGCGGAACCGGTGA
GTACACCGGAATTGCCAGGACGACCGGGTCCTTTCTTGGATTAACCCGCTCAATGCCTGGAG
ATTTGGGCGTGCCCCCGCGAGACTGCTAGCCGAGTAGTGTTGGGTCGCGAAAGGCCTTGTGG
TACTGCCTGATAGGGTGCTTGCGAGTGCCCCGGGAGGTCTCGTAGACCGTGCATCATGAGCA
CAAATCCT

*MYC* 5’-UTR
GACCCCCGAGCTGTGCTGCTCGCGGCCGCCACCGCCGGGCCCCGGCCGTCCCTGGCTCCCCT
CCTGCCTCGAGAAGGGCAGGGCTTCTCAGAGGCTTGGCGGGAAAAAGAACGGAGGGAGGG
ATCGCGCTGAGTATAAAAGCCGGTTTTCGGGGCTTTATCTAACTCGCTGTAGTAATTCCAGC
GAGAGGCAGAGGGAGCGAGCGGGCGGCCGGCTAGGGTGGAAGAGCCGGGCGAGCAGAGCT
GCGCTGCGGGCGTCCTGGGAAGGGAGATCCGGAGCGAATAGGGGGCTTCGCCTCTGGCCCA
GCCCTCCCGCTGATCCCCCAGCCAGCGGTCCGCAACCCTTGCCGCATCCACGAAACTTTGCC
CATAGCAGCGGGCGGGCACTTTGCACTGGAACTTACAACACCCGAGCAAGGACGCGACTCT
CCCGACGCGGGGAGGCTATTCTGCCCATTTGGGGACACTTCCCCGCCGCTGCCAGGACCCGC
TTCTCTGAAAGGCTCTCCTTGCAGCTGCTTAGACG

*ATF4* 5’-UTR (red ATG, uORF start codon)
TTTCTACTTTGCCCGCCCACAGATGTAGTTTTCTCTGCGCGTGTGCGTTTCCCTCCTCCCCGC
CCTCAGGGTCCACGGCCACCATGGCGTATTAGGGGCAGCAGTGCCTGCGGCAGCATTGGCCT
TTGCAGCGGCGGCAGCAGCACCAGGCTCTGCAGCGGCAACCCCCAGCGGCTTAAGCCATGG
CGCTTCTCACGGCATTCAGCAGCAGCGTTGCTGTAACCGACAAAGACACCTTCGAATTAAGC
ACATTCCTCGATTCCAGCAAAGCACCGCAACA
*ATF4* ΔuORF1 – first ATG mutated to AGG
*ATF4* ΔuORF2 – second ATG mutated to AGG

*HBB* 5’-UTR
ACATTTGCTTCTGACACAACTGTGTTCACTAGCAACCTCAAACAGACACC

### SUPPLEMENTARY FIGURES

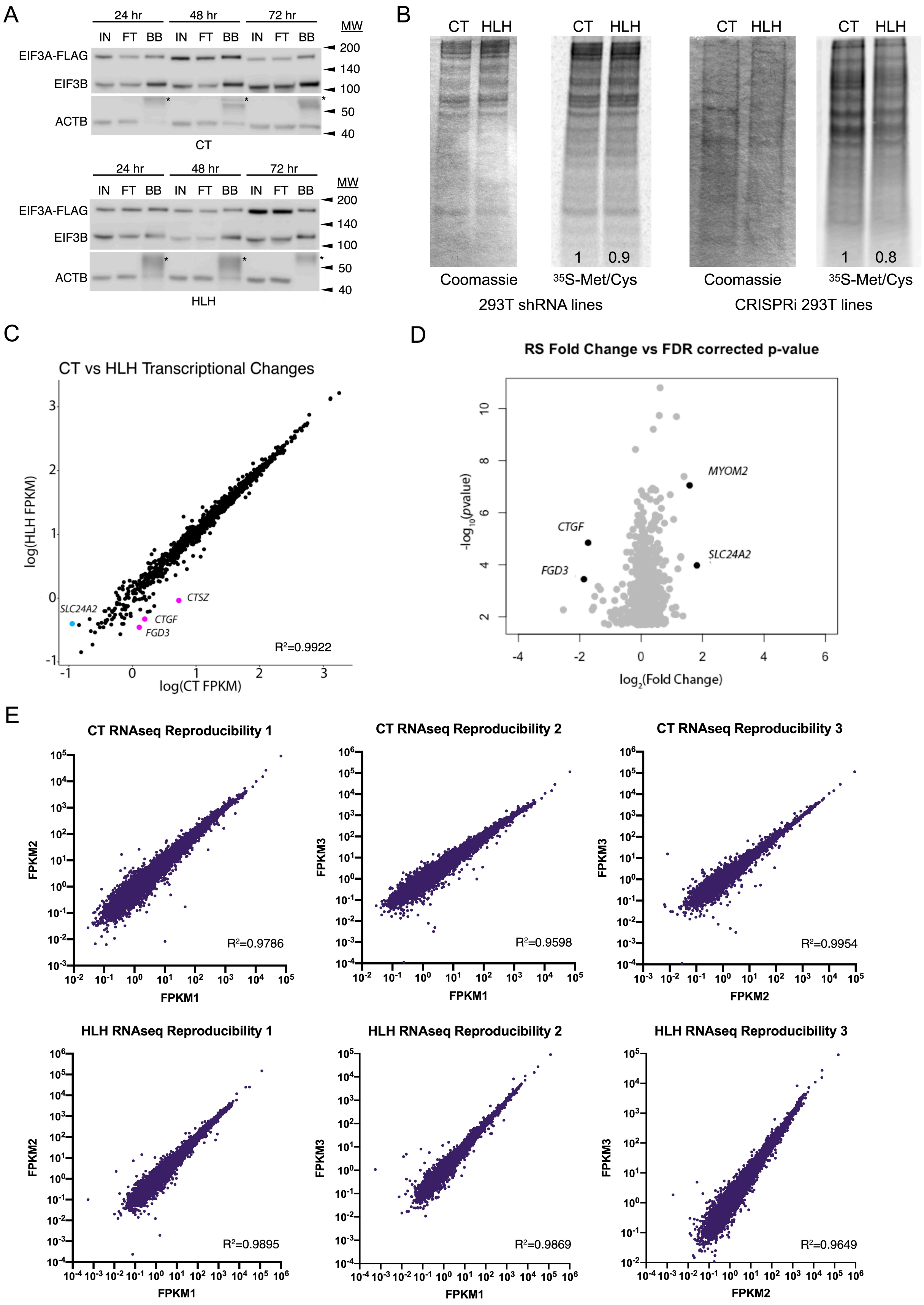

A
24 hr 48 hr 72 hr
IN FT BB IN FT BB IN FT BB
MW
EIF3A-FLAG
EIF3B
ACTB
200 140 100 50 40
CT
HLH
B
CT HLH
1 0.9
1 0.8
Coomassie
35S-Met/Cys
293T shRNA lines
CRISPRi 293T lines
C
CT vs HLH Transcriptional Changes
log(HLH FPKM)
log(CT FPKM)
SLC24A2
CTSZ
CTGF
FGD3
R²=0.9922
D
RS Fold Change vs FDR corrected p-value
MYOM2
CTGF
SLC24A2
FGD3
-log10(pvalue)
log2(Fold Change)
E
CT RNAseq Reproducibility 1
CT RNAseq Reproducibility 2
CT RNAseq Reproducibility 3
R²=0.9786
R²=0.9598
R²=0.9954
HLH RNAseq Reproducibility 1
HLH RNAseq Reproducibility 2
HLH RNAseq Reproducibility 3
R²=0.9895
R²=0.9869
R²=0.9649
FPKM1
FPKM2
FPKM3

**Fig. S1. Global translation and transcription analysis of eIF3A HLH* cell lines.** A. Transient transfection of FLAG-tagged CT and HLH* lentiviral vectors into HEK293T cells was analyzed by Western blotting for EIF3B and FLAG peptide. Levels of protein normalized to ACTB control. IN: input, FT: flowthrough, BB: bead bound. Asterisk indicates secondary antibody binding to EIF3B antibody used for pulldown. B. Representative $^{35}$S-Met/Cys metabolic labeling gels of the shRNA lentiviral HEK294T and CRISPRi cell lines. Total protein is stained by Coommassie (left). Numbers indicate fraction of $^{35}$S-Met/Cys signal normalized to total protein. C. Scatter plot of CT versus HLH* (HLH) transcriptional changes using average FPKM values of three biological replicates, showing transcripts meeting a *p*-value cutoff of 0.01. Three transcripts in pink are downregulated >3x, and one transcript in blue is upregulated >3x. D. Volcano plots of transcriptional fold change against FDR-corrected *p*-value, showing transcripts meeting an FDR cutoff of 0.05. Four transcripts in black show a >2x change in expression. E. RNAseq reproducibility scatter plots of CT and HLH biological replicates, showing non-zero transcripts used for statistical analysis. Highlighted in purple are transcripts with a >3x change in expression. $R^2$ coefficient of determination value is listed on the graph.

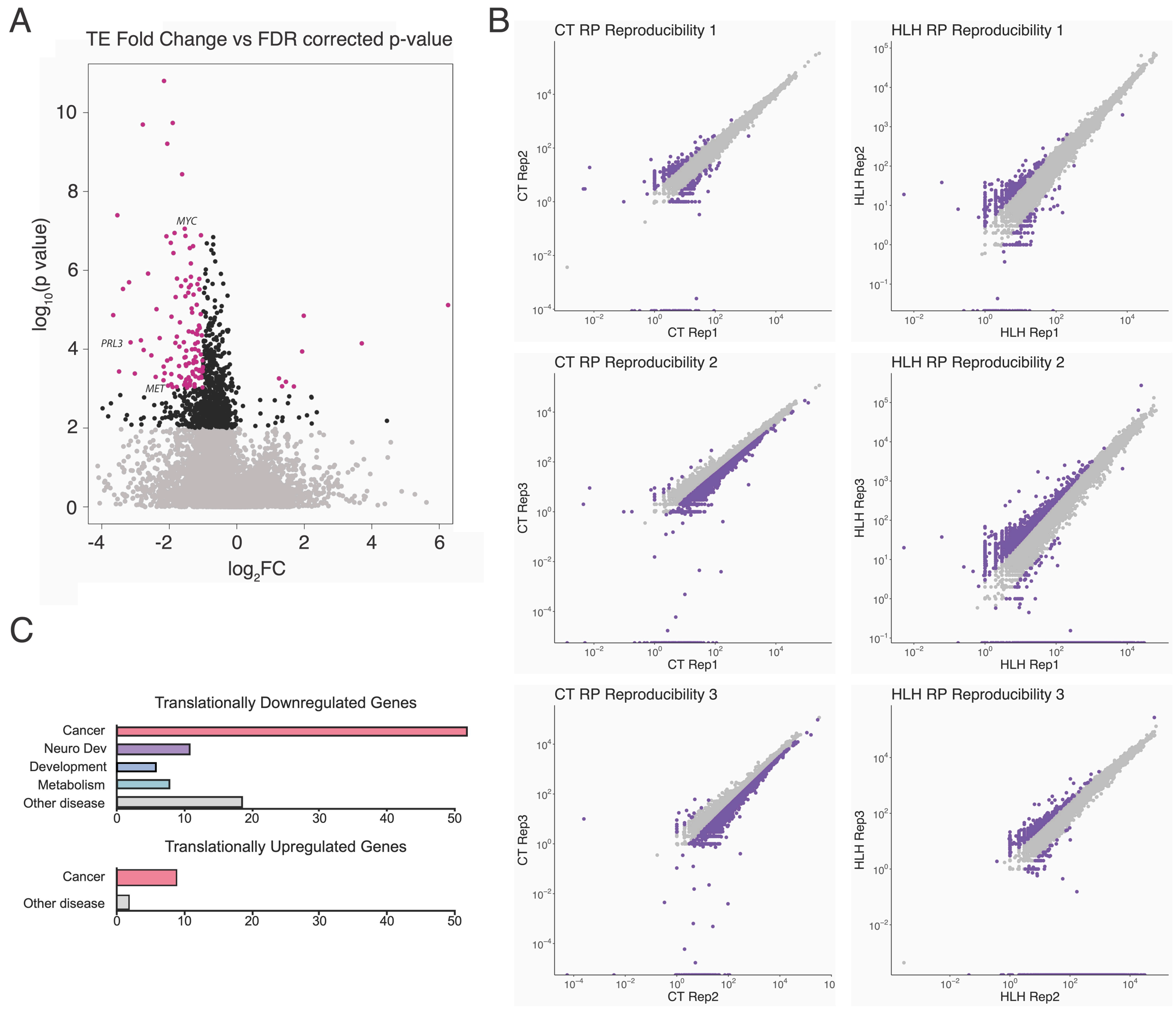


**Fig. S2. Downregulation of translation efficiency of proliferative mRNAs in HLH* cells.** A. Transcripts meeting an FDR corrected *p*-value cutoff of 0.05 in black. Out of those, transcripts with >3x fold change are highlighted in magenta. B. Ribosome profiling reproducibility scatter plots of CT and HLH* (HLH) biological replicates, showing non-zero transcripts used for statistical analysis. $R^2$ coefficient of determination value is listed on the graph. C. Functional classification of regulated transcripts based on literature analysis. See tables S5 and S6.

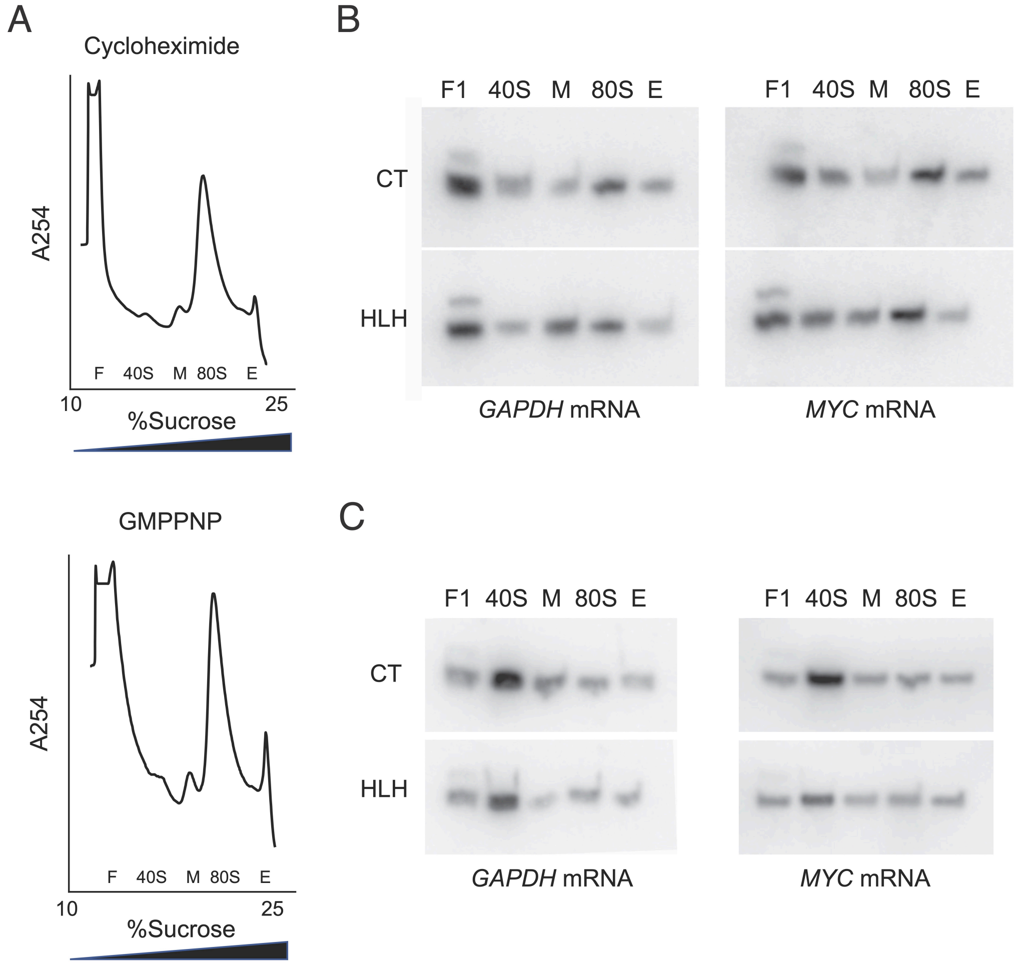


**Fig. S3. Met-tRNAi incorporation is unaffected by the eIF3A HLH mutation.** A. Sucrose gradient profiles of cycloheximide (top) and GMPPNP (bottom) stalled *in vitro* translation reactions fractionated on 10-25% sucrose gradients. B. Northern blotting of Met-tRNA$_i$ in the presence of cycloheximide. C. Northern blotting of tRNA$_i$ in the presence of GMPPNP.

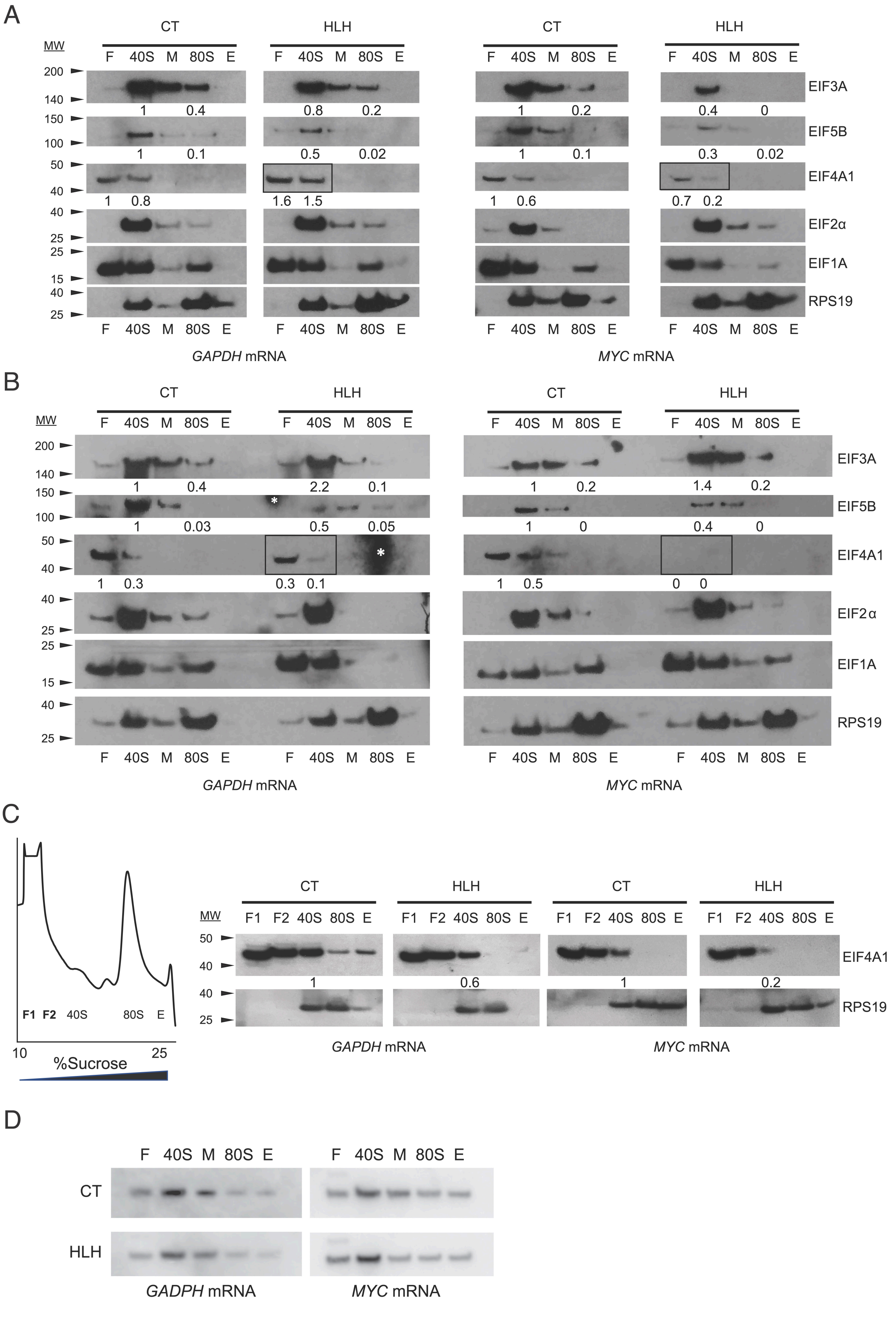
A
CT
HLH
CT
HLH
MW
F 40S M 80S E
200
140
150
100
50
40
40
25
25
15
40
25
EIF3A
1 0.4
0.8 0.2
1 0.2
0.4 0
EIF5B
1 0.1
0.5 0.02
1 0.1
0.3 0.02
EIF4A1
1 0.8
1.6 1.5
1 0.6
0.7 0.2
EIF2α
EIF1A
RPS19
GAPDH mRNA
MYC mRNA
B
EIF3A
1 0.4
2.2 0.1
1 0.2
1.4 0.2
EIF5B
1 0.03
0.5 0.05
1 0
0.4 0
EIF4A1
1 0.3
0.3 0.1
1 0.5
0 0
EIF2α
EIF1A
RPS19
GAPDH mRNA
MYC mRNA
C
F1 F2 40S 80S E
10
25
%Sucrose
F1 F2 40S 80S E
EIF4A1
1
0.6
1
0.2
RPS19
GAPDH mRNA
MYC mRNA
D
F 40S M 80S E
CT
HLH
GADPH mRNA
MYC mRNA

**Fig. S4. eIF3A HLH mutation causes transcript-specific defects in initiation factor eIF4A1 recruitment.** A. Western blot analysis of initiation factors in GMPPNP stalled *in vitro* translation reactions resolved by sucrose gradient fractionation, expanded from Fig. 2 to include additional factors. B. Western blot analysis of initiation factors in GMPPNP/RocA stalled *in vitro* translation reactions, expanded from Fig. 3 to include additional initiation factors. Boxes indicate fractions of interest for EIF4A1 levels. Asterisk indicates background signal in gel, which does not interfere with initiation factor distribution analysis. C. Western blot analysis of initiation factor EIF4A1 in GMPPNP/RocA stalled *in vitro* translation reactions resolved by sucrose gradient fractionation, expanded to include the top of gradient fractions F1 and F2. Note that the different lot of antibody used here was significantly more sensitive than in panels A and B. D. Northern blot analysis of Met-$tRNA_i$ distribution in GMPPNP/RocA stalled *in vitro* translation reactions.

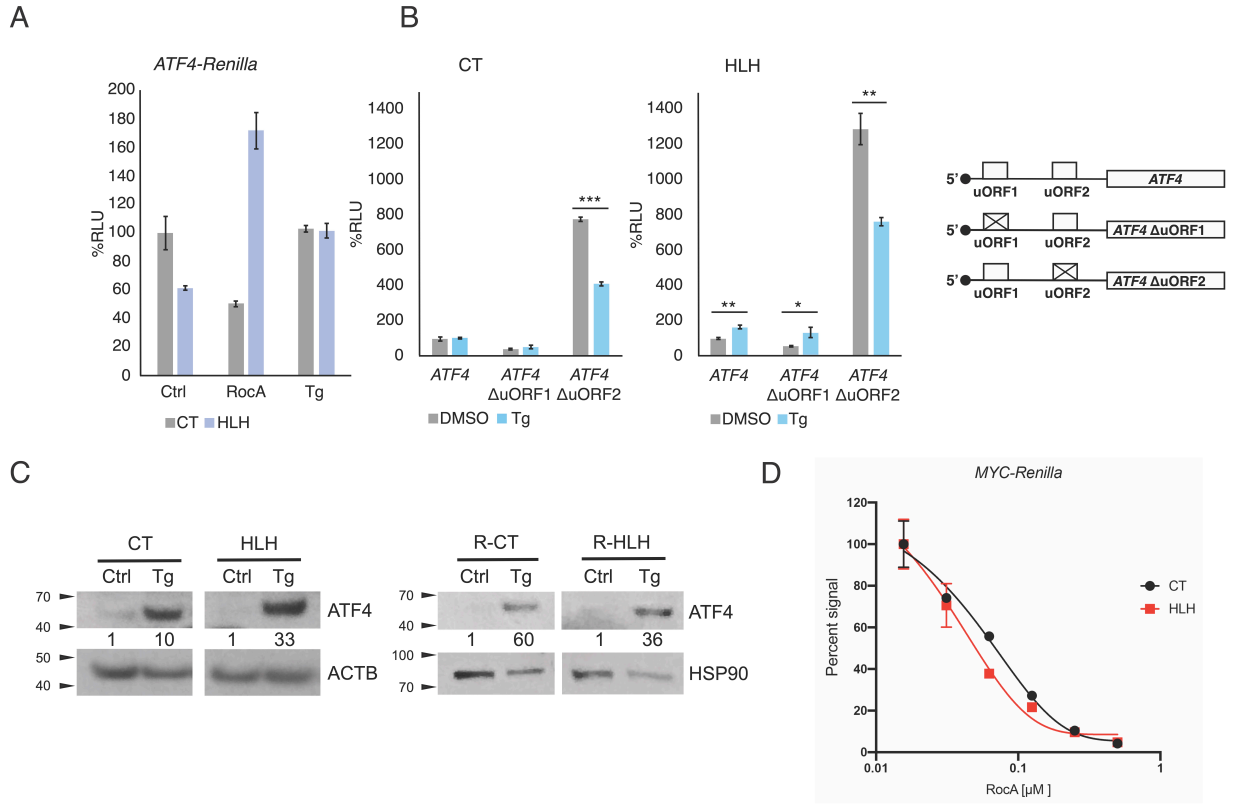


**Fig S5. eIF3A HLH mutation alters uORF-regulated translation in response to stress.** A. Live cell transfections of mRNAs containing the *ATF4* 5’-UTR fused to the *Renilla* luciferase ORF in the presence of thapsigargin and RocA. Relative Luciferase Units (RLU) percentage was normalized to internal *HBB* 5’-UTR-*Firefly* luciferase mRNA control signal. B. Live cell transfections of *ATF4* uORF variant 5’-UTRs (WT, ΔuORF1, ΔuORF2) fused to *Renilla* mRNAs in the presence of thapsigargin. Control samples are identical to those plotted in Fig. 3F. C. CRISPRi 293T cell lines (CT, HLH) and Ramos cell lines (R-CT, R-HLH) show induction of endogenous ATF4 upon thapsigargin treatment. D. Live cell transfection of *MYC* 5’-UTR – *Renilla* luciferase mRNA was performed to observe the effect of RocA-induced stress. Relative Luciferase Units (RLU) percentage was normalized to an internal *HBB* 5’-UTR-*Firefly* luciferase control signal.

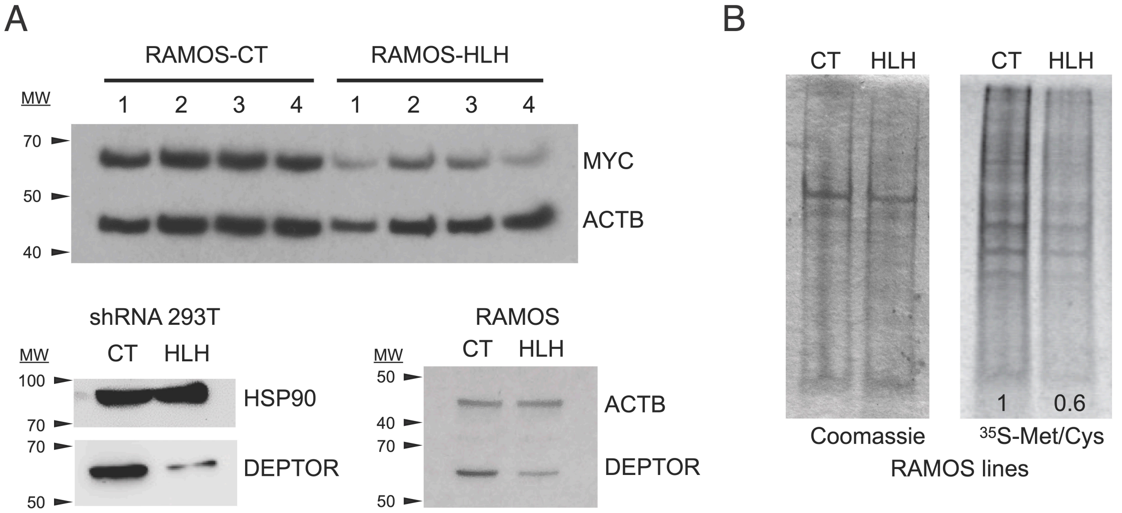


**Fig. S6. eIF3A HLH mutation leads to global translation decrease and loss of MYC in Burkitt's lymphoma cell lines.** A. Western blot validation of MYC suppression in Ramos HLH* cells. The numbers 1-4 represent separate cell lines transduced in parallel. Bottom panels show Western blot validation of additional cancer-associated negatively regulated transcript *DEPTOR* in HEK293T shRNA and Ramos shRNA cell lines. Levels of protein normalized to ACTB or HSP90 control given below gels. B. Representative $^{35}$S-Met/Cys metabolic labeling gels of the CRISPRi and Ramos cell lines. Total protein is stained by Coomassie (left). Numbers indicate fraction of $^{35}$S-Met/Cys signal normalized to total protein.
